# Longitudinal Metabolomic Profiling of Biogenic Amines in Plasma and CSF, and Their Correlation, Reveals Sex-Specific and Age Changes in TgF344 Alzheimer’s Disease Transgenic and Wildtype Rats

**DOI:** 10.64898/2025.12.03.692185

**Authors:** Chunyuan Yin, Imke Nelen, Amy Harms, Robin Hartman, Sabine Bos, Charlotte Nigh-van Kooij, Thomas Hankemeier, Alida Kindt, Elizabeth de Lange

## Abstract

**Background:** Alterations in amine metabolism have been implicated Alzheimer’s disease (AD). Cerebrospinal fluid (CSF) and plasma are key biofluids in AD research. CSF is considered to better reflect brain metabolic alterations than plasma, while plasma can be obtained more easily. However, plasma-CSF relationships are unclear.

**Aim:** To investigate longitudinal changes of amines in plasma and CSF, and their correlation across the two, in male and female TgF344 AD transgenic versus wildtype (WT) rats.

**Method:** LC-MS-based targeted metabolomics was used to analyze 60 and 55 amines in plasma and CSF, respectively, of male and female TgF344-AD and WT rats, at 12, 25, 50 and 85 weeks. Statistical analysis was performed using generalized logistic regressions, Pearson correlations, and differential correlations between groups and matrixes.

**Results:** Compared to WT controls, at *12 weeks*, TgF344-AD rats showed an increase of 3-methylhistidine, anserine, cysteine, s-methylcysteine, while at *25 weeks*, male TgF344-AD rats showed pronounced increases in CSF levels of alpha-aminobutyric acid, asparagine, glycylglycine, glycylproline, histidine, isoleucine, kynurenine, leucine, methionine, methionine sulfone, norepinephrine, phenylalanine, proline, tyrosine, and valine. At *50 weeks*, female TgF344-AD rats exhibited reductions in CSF for DL-3-aminoisobutyric acid, gamma-aminobutyric acid, ornithine, and putrescine. Distinct plasma-CSF correlations were found for 1-methylhistidine, 2-aminoadipic acid, putrescine, kynurenine, N6,N6,N6-trimethyl-lysine, DL-3-aminoisobutyric acid, and taurine, particularly in TgF344-AD rats.

**Conclusions:** Body fluid, age- and sex-dependent amine alterations in CSF and plasma of TgF344-AD rats compared to WT controls provide important insights into AD disease processes and may aid early diagnosis and therapeutic targeting.

## 1. Introduction

Alzheimer’s disease (AD) is the leading cause of dementia worldwide, characterized by progressive memory loss and cognitive decline [1]. Its hallmark pathological features include the accumulation of amyloid-beta plaques in the extracellular space and the formation of neurofibrillary tangles composed of hyperphosphorylated tau within neurons [2, 3]. These processes result in neuronal damage, synaptic dysfunction, and brain atrophy, which begin years to decades before clinical symptoms emerge [4]. While recent therapeutic advancements, such as the FDA-approved Aduhelm (aducanumab) and Leqembi (lecanemab), target amyloid pathology, their effectiveness is largely restricted to early disease stages [5, 6]. This underscores the urgent need for early biomarkers and a deeper understanding of AD-related metabolic alterations to improve diagnosis and therapeutic strategies, particularly for later disease stages where current treatments offer limited benefits.

Biogenic amines play important roles in various physiological processes, including acting as neurotransmitters, e.g., dopamine, glutamate, or serotonin [7], regulating blood pressure, e.g., histamine or tyramine, and contributing to cellular signaling and metabolic pathways, all of which are vital for maintaining brain function and systemic homeostasis [8, 9]. Alterations in amine metabolism have been increasingly implicated in neurodegenerative diseases, particularly AD [10, 11]. For instance, the amino acid glutamate is a key excitatory neurotransmitter and essential for synaptic transmission, but its dysregulation can lead to excitotoxicity, a process causing neuronal death, which is characteristic of AD progression [12, 13]. Gamma-aminobutyric acid (GABA), an inhibitory neurotransmitter, is also disrupted in AD, contributing to cognitive deficits and neuronal dysfunction [14]. In addition to neurotransmission, branched-chain amino acids (BCAAs) are involved in maintaining brain homeostasis, and disturbances in their metabolism may exacerbate neurodegeneration [15]. Evidence from both human and animal studies suggests that altered levels of amines such as serine, glycine, and alanine in cerebrospinal fluid (CSF) and plasma are associated with both early and late stages of AD, making them promising candidates for biomarkers of early disease progresses [16-18].

Direct brain sampling is only feasible postmortem, while CSF is the body-fluid in closest contact with the brain as it could potentially reflect brain-specific metabolic and pathological changes the most. However, its invasive collection limits widespread clinical use [19]. Plasma, although being more accessible and cost-effective in sampling, mainly reflects systemic metabolism and peripheral processes, but may partly reflect central processes. Therefore, plasma metabolomic profiling could serve as a valuable tool for tracking both systemic and brain-related metabolic disturbances during AD progression. Recent metabolomics studies have shown that certain metabolic changes in plasma closely mirror those in CSF, including disturbances in amino acid and polyamine metabolism, lysine degradation, and energy pathways [16, 20]. These findings highlight the potential of plasma profiling to reflect central metabolic alterations, supporting its role as a complementary biofluid for biomarker discovery. However, so far, only few human studies have addressed changes in amine profiles in the CSF and plasma in AD [16, 21], and more information is needed to understand the role of amines in AD progression in relation to their potential alterations in both fluids. As AD progression is closely associated with ageing, longitudinal studies should be performed comparing AD to controls, also addressing sex/gender dependencies [22].

Direct plasma-CSF correlations could support the use of plasma amines as potential non-invasive biomarkers for AD-related metabolic alterations. In our previous work we characterized age-, sex-, and AD-specific lipidomic alterations in plasma from TgF344-AD rats [22]. In this study, we explored CSF and plasma amine alterations of the same male and female TgF344-AD rats and their wild-type (WT) littermates at 12, 25, 50 and 85 weeks. Using targeted UPLC-MS we quantified amines in plasma and CSF, and compared their levels in the different age groups, sex and across biofluids in WT vs transgenic AD rats.

## 2. Materials and methods

### 2.1 Animals and sample collection

TgF344-AD rats and their age-matched wild-type (WT) Fischer F344 littermates were obtained and bred as previously described [22]. Briefly, this longitudinal study included 75 TgF344-AD rats and 74 WT littermates of both sexes, sampled at 12, 25, 50, and 85 weeks (detailed in Table 1). Housing conditions, including 12h/12h light-dark cycles, controlled temperature (21±2°C), humidity (40-60%) and standard rodent diet, were maintained as previously reported. Animal protocols were approved by the Leiden University Animal Welfare Body (AWB: AVD1060020171766). Blood and CSF samples were collected following established procedures [22]. Blood was drawn from the left ventricle of the heart into ethylenediaminetetraacetic acid (EDTA) tubes, and CSF was aspirated directly from the Cisterna magna. Both sample types were centrifuged at 2300G for 10 minutes at 4°C, aliquoted, and stored at -80°C until further analysis.

**Table 1.** Number of CSF and plasma samples for males and females at different ages in the WT and TgF344-AD rat groups; w = weeks.

| Age |  | WT rats |  |  |  | TgF344-AD rats |  |  |  |
| --- | --- | --- | --- | --- | --- | --- | --- | --- | --- |
|  |  | 12 w | 25 w | 50 w | 85 w | 12 w | 25 w | 50 w | 85 w |
| CSF | female | 5 | 7 | 8 | 9 | 6 | 10 | 10 | 11 |
| Plasma | male | 10 | 11 | 8 | 9 | 7 | 9 | 8 | 7 |
|  | female | 7 | 8 | 8 | 11 | 8 | 10 | 10 | 12 |
|  | male | 9 | 12 | 9 | 10 | 9 | 9 | 9 | 8 |

### 2.2 Amine measurement

Plasma and CSF samples were analyzed on the amine platform at the Biomedical Metabolomics Facility Leiden (BMFL), which quantifies biogenic amines using an AccQ-Tag derivatization method, adapted from the Waters protocol as previously described [23].

For each analysis, 5.0 μL of plasma or 10 μL of CSF was mixed with 5.0 μL of a spiked internal standard solution. Proteins were then precipitated by the addition of methanol, after which the samples were dried in a SpeedVac (Thermo). The residue was reconstituted in a borate buffer (pH 8.8) with AQC reagent. After the reaction, the samples were acidified with 10 μL of 20% formic acid and transferred to autosampler vials, then placed in an autosampler tray and cooled to 4°C until injection. Finally, 1.0 μL of the reaction mixture was injected into the UPLC-MS/MS system for targeted quantification of amines. Chromatographic separation was performed using an Agilent 1290 Infinity II LC System equipped with an AccQ-Tag Ultra column (Waters), operating at a flow rate of 0.7 mL/min over an 11-minute gradient. The UPLC was coupled to a triple quadrupole mass spectrometer (AB SCIEX Qtrap 6500) utilizing electrospray ionization. Analytes were detected in the positive ion mode and monitored using Multiple Reaction Monitoring (MRM) with nominal mass resolution. The acquired data were evaluated using MultiQuant Software for Quantitative Analysis (AB SCIEX, Version 3.0.2), by integration of assigned MRM peaks and normalization using the accordingly selected internal standards. For amino acid analysis, 13C15N-labeled analogs were utilized. For other amines, the closest-eluting internal standard was employed. Blank samples were used to assess background signal. Batch correction was performed using median normalization of the pooled quality control samples [24]. A total of 55 amines features from CSF and 60 from plasma with relative standard deviation of the quality control samples <30% passed the quality evaluation and were included in the statistical analysis.

### 2.3. Statistical analysis

All statistical tests were performed in R Studio (Version 2024.9.0.375). Amines passing quality evaluation were log2 transformed before analysis to normalize data distribution and reduce heterogeneity of variance.

#### Generalized logistic regression model

Univariate differences of amines in CSF and plasma between TgF344-AD and age-matched WT rats were assessed using a generalized logistic regression model. Analyses were conducted for both overall comparisons and subgroup analyses, controlling for sex and age (12, 25, 50, and 85 weeks). Metabolite estimate values and p-values across different groups were extracted from the summary of the models. P-values were adjusted for multiple comparisons using the Benjamini-Hochberg method to control the false discovery rate (FDR), and metabolites with FDR-adjusted q-value < 0.2 were considered statistically significant. All plots were generated using the `ggplot2` package in R.

#### Correlation analysis

To investigate the relationships between CSF and plasma metabolites, Pearson correlation coefficients were calculated using “cor.test” function for each metabolite across different disease groups (WT and TgF344-AD). Correlation analyses were classed by age (12, 25, 50, and 85 weeks) and sex (male and female) to identify age- and sex-specific patterns. Significant correlations were determined based on |R| ≥ 0.35, with strong correlations defined as |R| ≥ 0.6 and moderate correlations as 0.35 ≤ |R| < 0.6. Correlation heatmaps were generated using the `pheatmap` package in R, with hierarchical clustering applied to highlight metabolite groups exhibiting similar correlation patterns. Correlations falling within the range of -0.35 to 0.35 were considered weak and marked as blank to improve visualization. Grouping in the heatmap is by both sex (male and female) and age (12, 25, 50, and 85 weeks), allowing for the exploration of sex- and age-dependent correlation patterns between CSF and plasma metabolites. Comparisons were performed separately for WT and TgF344-AD rats to assess disease-specific correlation differences across sex and age. Differential correlation analysis was performed using Fisher’s r-to-z transformation to generate a z-score for the difference between AD and WT correlation coefficients for each metabolite in general and separated by gender and age.

## 3. Results

### 3.1 Age- and sex-specific alterations in CSF and plasma amines profiles in TgF344-AD and wildtype rats

#### Alterations in CSF biogenic amine profiles

CSF amine differences between TgF344-AD rats and age-matched WT rats, assessed by generalized logistic regression models, are summarized in **Figure 1**. The figure illustrates estimate group effects (β coefficients) across sex and age, where orange indicates higher levels in TgF344-AD rats and purple indicates lower levels compared to WT rats.

**Figure 1.**
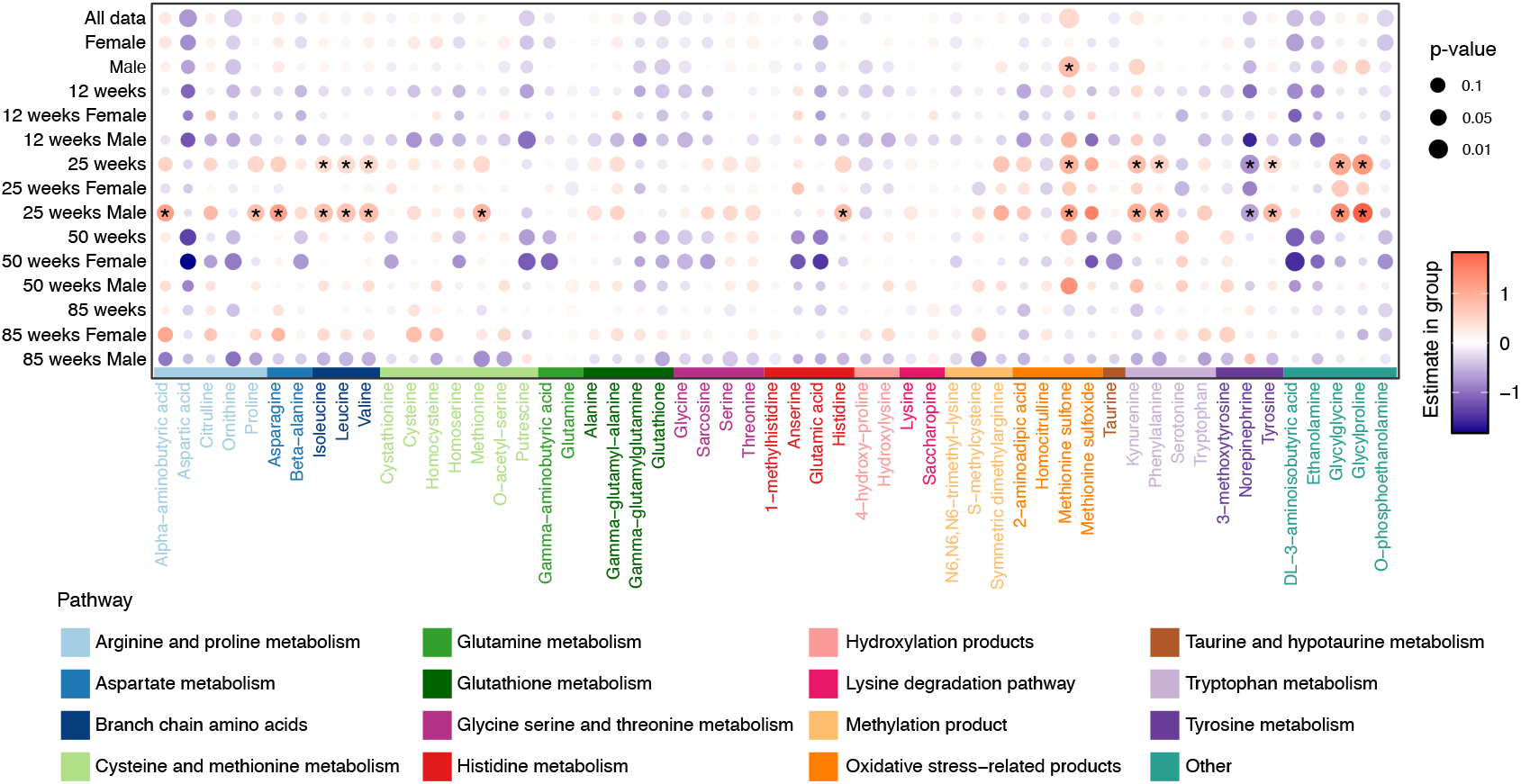
Estimated differences in CSF amines comparing TgF344-AD with age-matched wildtype (WT) rats based on generalized logistic regression models (GLMs). Each dot represents the estimated group difference (β coefficient), with color indicating the direction of change (orange = higher in TgF344-AD; purple = lower) and dot size reflecting the significance level. Metabolites passing FDR correction (q < 0.2) are marked with an asterisk (*). Biogenic amines are color-coded according to their associated metabolic pathways, such as arginine and proline metabolism, glutathione metabolism, or oxidative stress-related products, as indicated in the legend. Corresponding p-values and FDR-adjusted q-values for all tested comparisons are provided in **Supplementary Table-1**.

Overall, the analysis revealed clear age- and sex-dependent alterations in CSF biogenic amines across disease progression. At *12 weeks*, changes were minimal, with only putrescine showing downregulated in male TgF344-AD rats compared WT rats, while no metabolite changes were observed in female rats. By *25 weeks*, male TgF344-AD rats exhibited a broad increase in multiple biogenic amine and related metabolites (e.g., alpha-aminobutyric acid, asparagine, glycylglycine, glycylproline, histidine, isoleucine, kynurenine, leucine, methionine, methionine sulfone, phenylalanine, proline, tyrosine, and valine). A marked decrease was found for norepinephrine. In contrast, at 50 weeks, female TgF344-AD rats displayed pronounced decreases in several amines, including DL-3-aminoisobutyric acid, gamma-aminobutyric acid, ornithine, and putrescine, while 50-week male TgF344-AD rats showed an increase only in methionine sulfone compared to 50-week male rats. At *85 weeks*, no CSF amines passed *p* < 0.05 between TgF344-AD and WT rats.

Together, these results indicate that male TgF344-AD rats exhibited early and widespread increases in CSF amine levels at 25 weeks, whereas females exhibit more pronounced decreases at 50 weeks, underscoring age-related and sex-associated dynamics during AD progression.

#### Alterations in plasma biogenic amine profiles

Plasma amine differences between TgF344-AD rats and age-matched WT rats are summarized in **Figure 2**. Like the CSF data, age- and sex-dependent metabolic alterations were evident in plasma.

**Figure 2.**
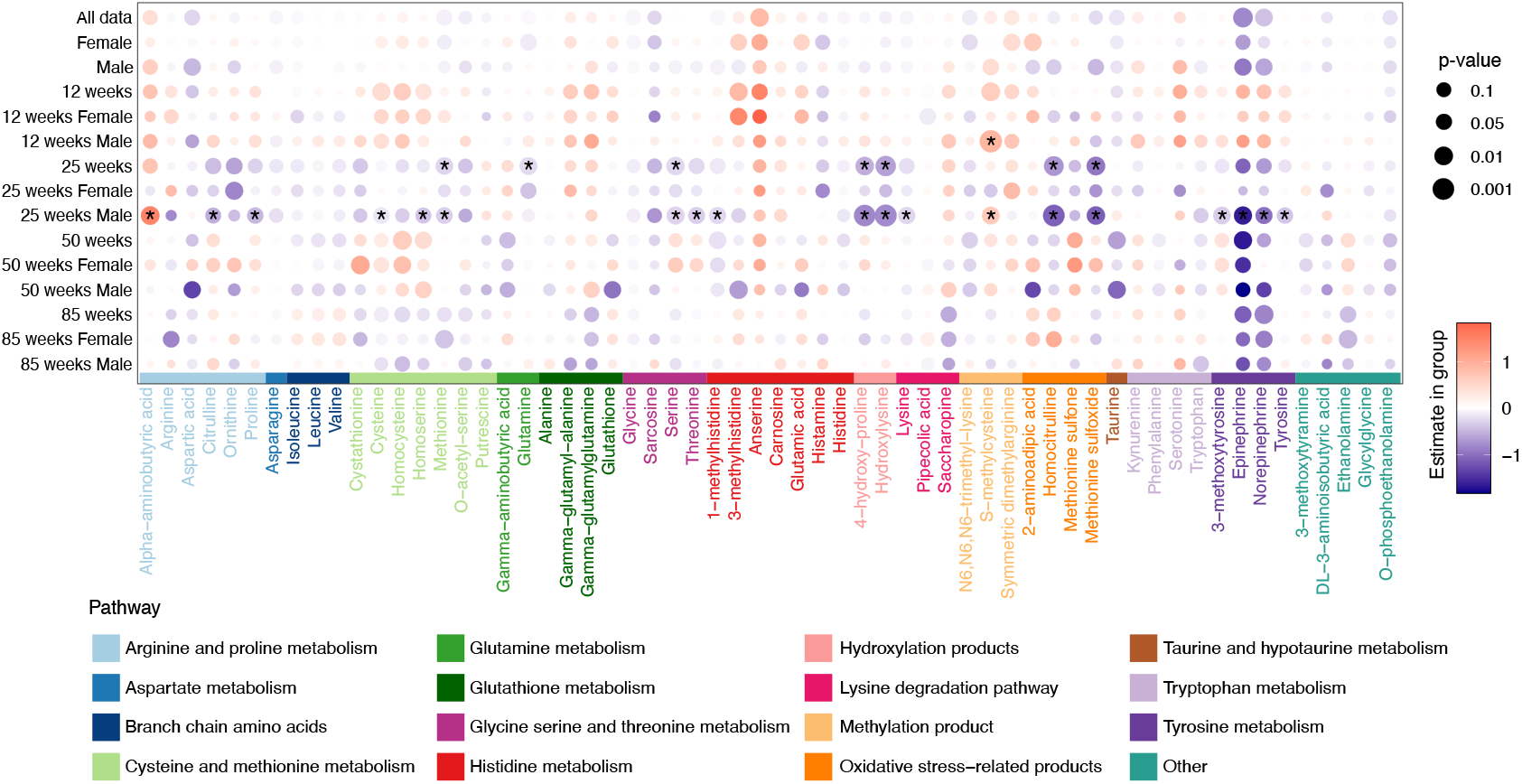
Estimated differences in plasma amines comparing TgF344-AD with age-matched wildtype (WT) rats based on generalized logistic regression models (GLMs). Each dot represents the estimated group difference (β coefficient), with color indicating the direction of change (red = higher in TgF344-AD; blue = lower) and dot size reflecting the significance level. Metabolites passing FDR correction (q < 0.2) are marked with an asterisk (*). Biogenic amines are color-coded according to their associated metabolic pathways, such as arginine and proline metabolism, glutathione metabolism, or oxidative stress-related products, as indicated in the legend. Corresponding p-values and FDR-adjusted q-values for all tested comparisons are provided in **Supplementary Table-2**.

At *12 weeks*, TgF344-AD rats showed increased levels of 3-methylhistidine, anserine, cysteine, homocysteine, homoserine, and s-methylcysteine compared to WT rats. At *25 weeks*, male TgF344-AD rats displayed widespread decreases in 1-methylhistidine, 3-methoxytyrosine, 4-hydroxy-proline, cysteine, epinephrine, homocitrulline, hydroxylysine, lysine, methionine, methionine sulfoxide, norepinephrine, proline, serine, threonine, tyrosine, while alpha-aminobutyric acid and s-methylcysteine were elevated. These findings were consistent with those in the 25-week combined gender groups, indicating that male-driven changes dominate at 25 weeks. At *50 weeks*, males TgF344-AD rats continued to show reduction in 3-methylhistidine, aspartic acid, glutathione, and taurine, while showing increases in gamma-glutamylglutamine and homoserine. At *85 weeks*, female TgF344-AD rats displayed decreased levels of arginine, ethanolamine, methionine, and norepinephrine.

Altogether, these results indicate that plasma amine alterations evolve dynamically with age and sex, showing early increases in young TgF344-AD rats and progressive declines at later stages, particular in males. These changes mirror the disease stage- and sex-dependent patterns observed in CSF, reflecting systemic manifestations of amine dysregulation during AD progression.

### 3.2 Pearson correlation between CSF and plasma biogenic amines in TgF344-AD and WT rats

CSF-plasma correlations across TgF344-AD and WT rats were summarized **in Figure 3**, which highlights the overall strength and direction of associations between the two biofluids. Of the 53 amines detected in both biofluids, 12 exhibited statistically significant (*q* < 0.05) CSF-plasma correlations in either TgF344-AD or WT rats. The *p*-values and FDR-adjusted *q*-values for these correlations, together with the Fisher’s z-transformed correlation coefficients, are provided in **Table 2**.

**Table 2.**
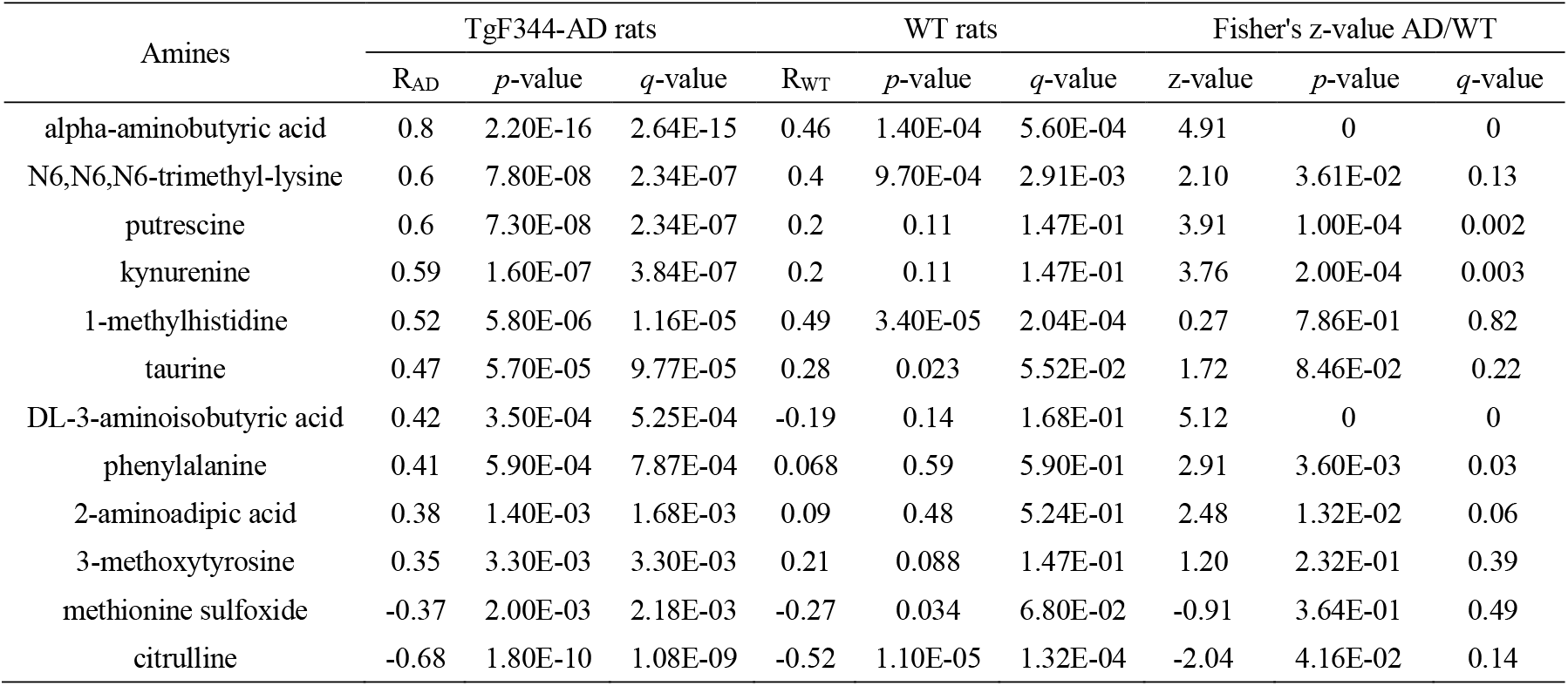
Top CSF-plasma correlated amines in TgF344-AD and WT rats. Amines are ranked based on the strength of their Pearson correlation coefficients (R) between CSF and plasma levels. For each amine, the correlation coefficient (R), associated *p*-values, and FDR-adjusted *q*-values are provided separately for TgF344-AD and WT rats. Positive R values indicate positive correlation between plasma and CSF concentrations of the amines. The Fisher’s z-value represent the statistical difference in correlation strength between TgF344-AD and WT rats. The amines are ranked ordered for the p-value in AD rats.

| Amines | TgF344-AD rats |  |  | WT rats |  |  | Fisher's z-value AD/WT |  |  |
| --- | --- | --- | --- | --- | --- | --- | --- | --- | --- |
|  | R<sub>AD</sub> | *p*-value | *q*-value | R<sub>WT</sub> | *p*-value | *q*-value | z-value | *p*-value | *q*-value |
| alpha-aminobutyric acid | 0.8 | 2.20E-16 | 2.64E-15 | 0.46 | 1.40E-04 | 5.60E-04 | 4.91 | 0 | 0 |
| N6,N6,N6-trimethyl-lysine | 0.6 | 7.80E-08 | 2.34E-07 | 0.4 | 9.70E-04 | 2.91E-03 | 2.10 | 3.61E-02 | 0.13 |
| putrescine | 0.6 | 7.30E-08 | 2.34E-07 | 0.2 | 0.11 | 1.47E-01 | 3.91 | 1.00E-04 | 0.002 |
| kynurenine | 0.59 | 1.60E-07 | 3.84E-07 | 0.2 | 0.11 | 1.47E-01 | 3.76 | 2.00E-04 | 0.003 |
| 1-methylhistidine | 0.52 | 5.80E-06 | 1.16E-05 | 0.49 | 3.40E-05 | 2.04E-04 | 0.27 | 7.86E-01 | 0.82 |
| taurine | 0.47 | 5.70E-05 | 9.77E-05 | 0.28 | 0.023 | 5.52E-02 | 1.72 | 8.46E-02 | 0.22 |
| DL-3-aminoisobutyric acid | 0.42 | 3.50E-04 | 5.25E-04 | -0.19 | 0.14 | 1.68E-01 | 5.12 | 0 | 0 |
| phenylalanine | 0.41 | 5.90E-04 | 7.87E-04 | 0.068 | 0.59 | 5.90E-01 | 2.91 | 3.60E-03 | 0.03 |
| 2-aminoadipic acid | 0.38 | 1.40E-03 | 1.68E-03 | 0.09 | 0.48 | 5.24E-01 | 2.48 | 1.32E-02 | 0.06 |
| 3-methoxytyrosine | 0.35 | 3.30E-03 | 3.30E-03 | 0.21 | 0.088 | 1.47E-01 | 1.20 | 2.32E-01 | 0.39 |
| methionine sulfoxide | -0.37 | 2.00E-03 | 2.18E-03 | -0.27 | 0.034 | 6.80E-02 | -0.91 | 3.64E-01 | 0.49 |
| citrulline | -0.68 | 1.80E-10 | 1.08E-09 | -0.52 | 1.10E-05 | 1.32E-04 | -2.04 | 4.16E-02 | 0.14 |

**Figure 3.**
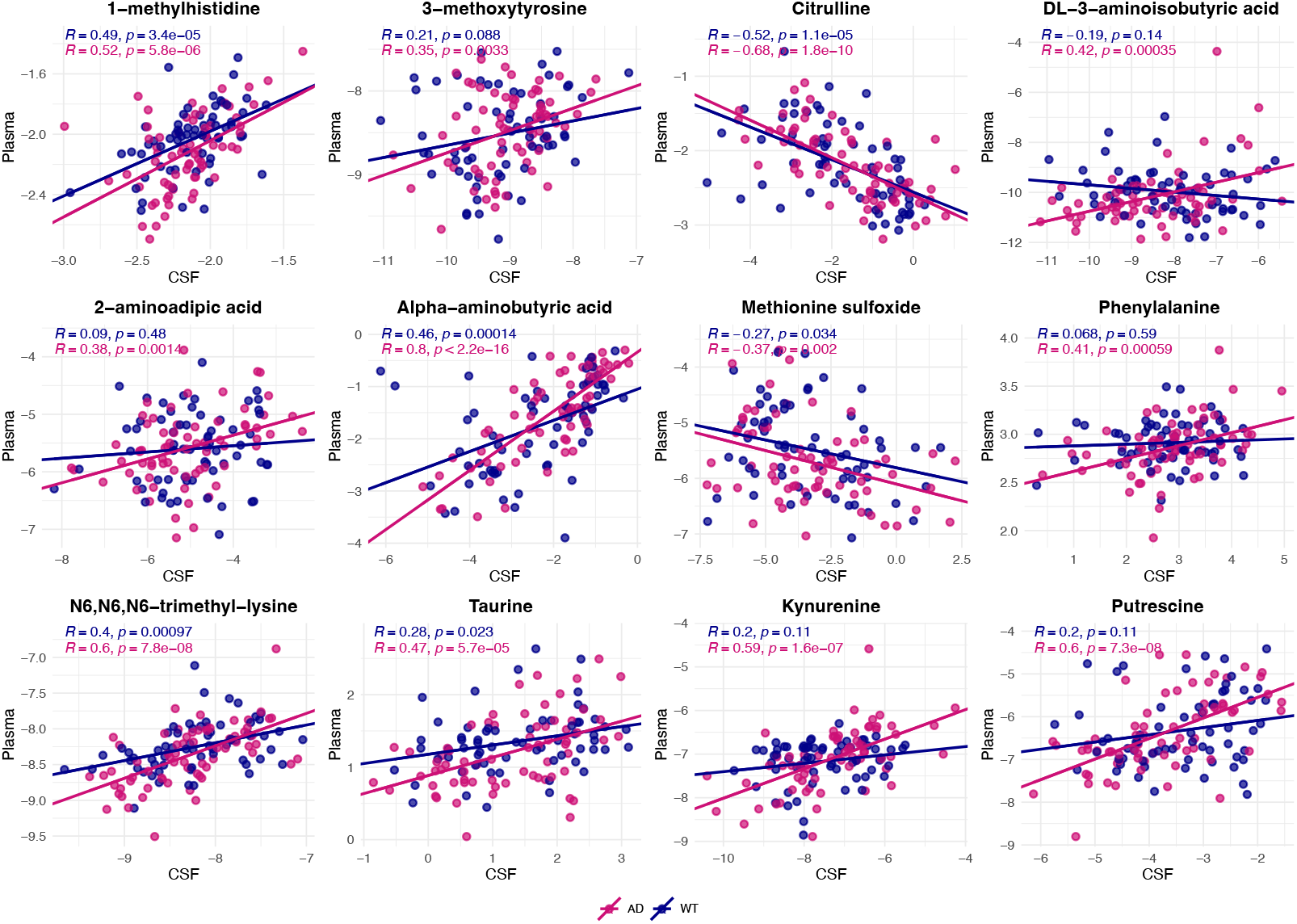
Pearson correlation analysis of CSF and plasma metabolites highlighting correlations within TgF344-AD (dark pink) and WT rats (purple). For each amine, a point represents the relative ratio of the amine measured in both CSF and plasma for one rat. Based on all points, a line with a correlation coefficient (R) that indicates the strength and direction of associations between CSF and plasma levels. In the TgF344-AD rats, strong correlations were observed for metabolites such as alpha-aminobutyric acid, citrulline, N6,N6,N6-trimethyl-lysine, and putrescine while the WT rats showed moderate correlations for metabolites like 1-methylhistidine, alpha-aminobutyric acid, citrulline, and N6,N6,N6-trimethyl-lysine. Strong correlations are defined as |R| ≥ 0.6, and moderate correlations as 0.4 ≤ |R| < 0.6.

Overall, TgF344-AD rats exhibited stronger CSF-plasma correlations than WT. Alpha-aminobutyric acid, citrulline, N6,N6,N6-trimethyl-lysine, and putrescine showed strong correlations (|R| ≥ 0.6) in the TgF344-AD rats, while these correlations were only moderate (0.4 ≤ |R|< 0.6) in WT rats. When stratified by sex **(Supplementary Table 4-5**), female rats exhibited more pronounced differences in CSF-plasma correlations between TgF344-AD and WT groups, particularly for alpha-aminobutyric acid, citrulline, and N6,N6,N6-trimethyl-lysine. Across ages (**Supplementary Table 6-9**), correlation patterns in citrulline evolved dynamically: WT rats showed stronger negative correlation at 25 weeks, while TgF344-AD became clearly more negative at 50 weeks and 85 weeks.

Taken together, these data show that some amines, *e*.*g*., alpha-aminobutyric acid, N6,N6,N6-trimethyl-lysine, exhibit AD-specific strengthening of plasma-CSF associations, especially in females, whereas others, *e*.*g*., citrulline, display sex- and age-dependent shifts that can favor support either WT (25 weeks) or AD (50-85 weeks).

### 3.3 Age- and sex-dependent correlation patterns between CSF and plasma amines in TgF344-AD and WT rats

To visualize the correlations between plasma and CSF amines in TgF344-AD and WT rats across all four age groups (12, 25, 50, and 85 weeks), Pearson correlation coefficients were calculated and visualized as a clustered heatmap (**Figure 4**). The heatmap reveals distinct clustering patterns of amines according to genotype, age, and sex, highlighting the progressive and sex-dependent alterations in plasma-CSF associations.

**Figure 4.**
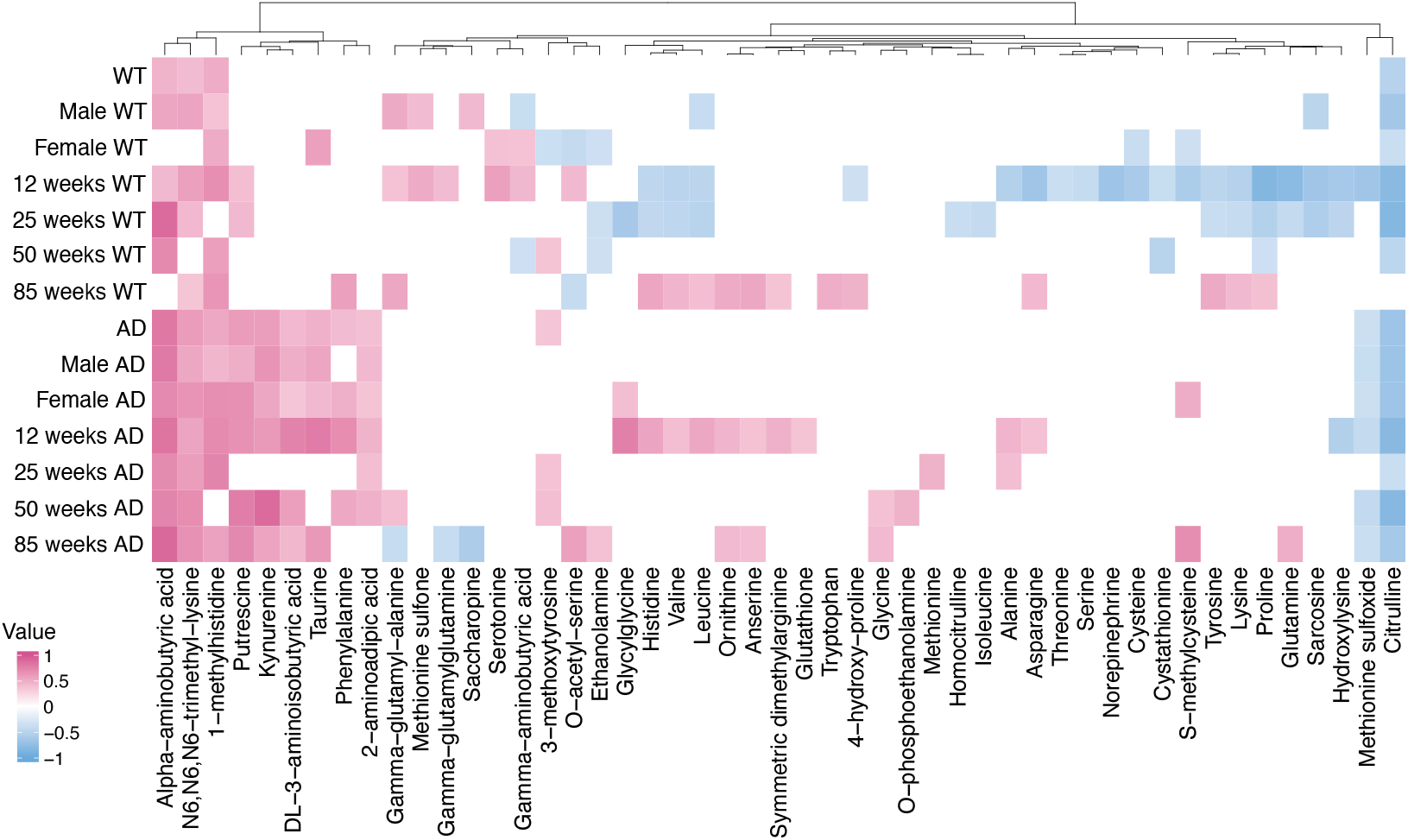
The heatmap illustrates the Pearson correlation results between metabolites in plasma and CSF across different age groups in TgF344-AD rats and WT rats. The plot highlights correlation patterns where positive correlations are shown in red and negative correlations in blue, while correlations with values between (-0.35)-0.35 were considered as too weak and blanked out. A full heatmap including all data is provided in **Supplementary Figure 1** for reference, and an alphabetically ordered version is shown in **Supplementary Figure 2**.

#### Disease-and age-dependent correlations

At *12 weeks*, TgF344-AD rats showed strong positive plasma-CSF correlations for several amines, including 2-aminoadipic acid, DL-3-aminoisobutyric acid, kynurenine, phenylalanine, putrescine, and taurine compared with WT rats. At *25 weeks*, TgF344-AD rats continued to exhibit strong positive correlations for 1-methylhistidine, alpha-aminobutyric acid, and N6,N6,N6-trimethyl-lysine. By *50 weeks*, positive correlations persisted for several amines such as alpha-aminobutyric acid, kynurenine, and putrescine in TgF344-AD rats. At *85 weeks*, WT rats exhibited stronger correlations for a different set of amines (e.g., 4-hydroxy-proline, asparagine, histidine, leucine, lysine, proline, symmetric dimethylarginine, tryptophan, tyrosine, and valine), while TgF344-AD rats showed weaker correlations.

#### Sex-specific plasma-CSF correlations

Distinct sex-dependent correlation patterns between TgF344-AD rats and WT rats were observed. In TgF344-AD rats, strong positive correlations (|R| ≥ 0.6) were found in females for metabolites such as glycylglycine, s-methylcysteine, and phenylalanine, which were absent in males. In WT rats, 3-methoxytyrosine, ethanolamine, o-acetyl-serine, serotonin, and taurine exhibited strong correlations in females but not in males, reflecting sex-linked metabolic coupling between plasma and CSF.

#### Cluster-based classification of correlation profiles

Based on the similarity of correlation profiles across genotype, age, and sex groups, three group trends were identified. The first group showed significant correlations in both WT and TgF344-AD rats but differed in their strength. This group included 1-methylhistidine, alpha-aminobutyric acid, and N6,N6,N6-trimethyl-lysine. The second group mainly showed correlations in the TgF344-AD rats, but not in the WT rats, and consisted of 2-aminoadipic acid, DL-3-aminoisobutyric acid, kynurenine, putrescine, phenylalanine, and taurine. The third group only showed negative correlations in WT rats at 12 weeks of age, and consisted of alanine, asparagine, citrulline, cysteine, cystathionine, glutamine, hydroxylysine, lysine, methionine sulfoxide, norepinephrine, proline, sarcosine, serine, s-methylcysteine, threonine, and tyrosine.

## Discussion

In this study, we investigated amine profiles in CSF and plasma samples from TgF344-AD and WT rats, assessing the influence of age and sex, as well as the relationships between the CSF and plasma amines. Of particular interest were the amine alterations occurring at early stages of AD progression. Male TgF344-AD rats predominantly exhibited amine changes at early disease stages (25 weeks), while female rats showed pronounced changes at a later stage (50 weeks). Additionally, correlations between plasma and CSF amine profiles were stronger in TgF344-AD rats compared to WT rats, suggesting potential alterations in blood-brain and/or blood-CSF barrier function under AD conditions. Together, these findings inform on AD related amine metabolism in a sex- and age-dependent manner, indicating the importance of explicitly addressing these factors, which is typically lacking in literature. The most prominent amine alterations are discussed in detail below.

### Early metabolic disruptions in male TgF344-AD rats involving oxidative stress, neuroinflammation, and neurotransmitter imbalance

The observed elevation of alpha-aminobutyric acid, asparagine, glycylproline, and methionine sulfone in the CSF of 25-week-old male TgF344-AD rats suggests potential metabolic disruptions associated with AD early-stage progression. Alpha-aminobutyric acid, linked to GABAergic systems [25], and asparagine, involved in nitrogen metabolism and neurotransmitter homeostasis [26, 27], suggest disruptions in neurotransmitter balance at early disease stages. Glycylproline, a dipeptide derived from collagen degradation, plays a role in neuroinflammation and blood-brain barrier (BBB) dysfunction in TgF344-AD rats [28]. Elevated glycylproline in male TgF344-AD rats may reflect increased neuroinflammatory responses, as AD is characterized by heightened neuroinflammation and microglial activation, particularly in male subjects [29]. Methionine sulfone, an oxidative product of methionine, accumulated in the CSF of male TgF344-AD rats, which may indicate heightened oxidative burden at 25 weeks, an early disease stage [30, 31]. This observation is consistent with previous human metabolomics findings, where oxidative stress related pathways, including those involving methionine sulfone, were found altered in both plasma and CSF of AD patients [20, 30, 32-34]. Further, the five metabolites associated with the transsulfuration pathway (cystathionine, cysteine, serine, s-methylcysteine, threonine) showed strong negative correlations across matrixes in 12-week-old WT rats but not in TgF344-AD rats. This pathway is critical for the maintenance of sulfur amino acid homeostasis with disruptions to this pathway being linked to neurodegenerative diseases, including AD [35, 36]. Given that the WT rats represent a healthy brain, an absence of these negative correlations could indicate early-stage disruptions leading.

Our findings support methionine sulfone as a sensitive indicator of oxidative imbalance in male TgF344-AD rats and are in line with prior studies reporting stronger oxidative stress signatures in males than in females in TgF344-AD model, potentially reflecting sex differences in antioxidant defense mechanisms [37].

### Sex-specific amine reductions in polyamine-related pathways at mid-stage Tgf344-AD in female rats

The observed decreases in DL-3-aminoisobutyric acid, gamma-aminobutyric acid, ornithine, and putrescine in the CSF of female TgF344-AD rats at 50 weeks suggest metabolic change distinct from that in males, indicating sex-specific metabolic disruptions associated with advanced-stage AD pathology. Notably, DL-3-aminoisobutyric acid (also known as β-aminoisobutyric acid) has been genetically linked to aging-related mild cognitive impairment, further supporting its relevance to neurodegenerative processes [38]. Ornithine and putrescine, key metabolites involved in polyamine metabolism and cellular stress responses [39-42], may reflect altered neuroprotective mechanisms or disrupted cellular homeostasis pathways [43, 44], potentially indicating increased vulnerability or impaired adaptive responses more pronounced in females at advanced stages of AD. Changes in polyamine metabolism have previously been associated with neuronal survival, regulation of oxidative stress, and modulation of neuroinflammation [39, 41], highlighting the importance of these amines in maintaining brain function during neurodegeneration.

### Enhanced amine plasma-CSF correlations in TgF344-AD rats suggest altered blood-brain barrier (BBB) transport

We observed strong plasma-CSF correlations for specific amines, including 1-methylhistidine, alpha-aminobutyric acid, N6,N6,N6-trimethyl-lysine, 2-aminoadipic acid, DL-3-aminoisobutyric acid, kynurenine, phenylalanine, putrescine, and taurine. Of these, the first three showed strong correlations between the two matrixes in all WT rats and all TgF3444-AD rats, regardless of age or sex, while the latter six are mainly correlating in the TgF344-AD rats. The enhanced plasma-CSF correlations of these amines in TgF344-AD rats could reflect altered permeability or transport across the BBB and blood-CSF barrier, potentially related to neuroinflammation, oxidative stress, and/or structural alterations associated with AD pathology [45-49]. Several of the amines mainly correlating in the TgF344-AD rats have known transport mechanisms across the BBB. For instance, 2-aminoadipic acid, an analog of lysine, is transported across the BBB via the system y+ and system L amino acid transporters [12, 50]. Kynurenine, produced via the tryptophan-kynurenine pathway, crosses the BBB predominantly through the L-type amino acid transporter 1 (LAT1), allowing kynurenine to enter the brain and contribute to neuroinflammation and neurodegeneration in AD [51-54]. Taurine, a neuroprotective and anti-inflammatory amino acid, may indicate adaptive or compensatory mechanisms responding to disease-associated stress [55-57].

Furthermore, Fisher’s r-to-z transformation analyses revealed significant differences in the strength of plasma-CSF amine correlations between TgF344-AD and WT groups. Key metabolites such as alpha-aminobutyric acid, N6,N6,N6-trimethyl-lysine, and citrulline showed differential correlation patterns in AD versus WT rats (**Supplementary Table 3**). Overall, alpha-aminobutyric acid and N6,N6,N6-trimethyl-lysine exhibited stronger positive plasma-CSF correlations in TgF344-AD rats than in WT, whereas citrulline became more negatively correlated in TgF344-AD rats. Sex- and age-specific patterns were also observed, with alpha-aminobutyric acid showing a significantly stronger positive correlation (q<0.2) in male TgF344-AD rats (Supplementary Table 4), and citrulline showing a weaker correlation in female TgF344-AD rats (Supplementary Table 5) when compared to their WT littermates. Across age, alpha-aminobutyric acid displayed a significant increase (q < 0.2) in correlation strength at 25 weeks, with WT rats showing a stronger plasma-CSF correlation than TgF344-AD rats (**Supplementary Table 7**), suggesting early disease-stage alterations in particular amine exchange between CSF and plasma.

The observed sex-specific differences in plasma-CSF amine associations highlight the necessity of incorporating sex as a critical variable in AD research. For example, certain amines such as glycylglycine, phenylalanine, and s-methylcysteine displayed strong plasma-CSF correlations in female TgF344-AD rats but not in male TgF344-AD rats, potentially due to sex-dependent regulation of amines in AD. Future studies should further investigate the mechanistic underpinnings of these amine shifts, particularly in relation to oxidative stress and neuroinflammation metabolism. Given the robust amine plasma-CSF associations observed, our findings highlight the potential utility of amines for monitoring AD progression and identifying disease-specific biomarkers.

### Advantages of the TgF344-AD model in translational studies

This TgF344-AD rat model offers unique opportunities for experimental interventions, including: (i) Tissue-Specific Pathophysiology Studies: Enabling a deeper investigation into brain-periphery metabolic interactions. (ii) Therapeutic Testing: Allowing for targeted modulation of metabolic pathways through pharmacological interventions. (iii) Mechanistic Validation: Providing opportunities to apply inhibitors or genetic modifications to dissect key disease-related pathways. Thus, our study not only underscores the amine changes associated with AD but also highlights the potential of the TgF344-AD rat model as a translational tool for mechanistic investigations and therapeutic advancements in AD research, as several of the here observed results were also observed in humans.

### Limitations of the study

This study exclusively analyzed amines, thereby capturing only a subset of the broader metabolic alterations associated with AD progression and aging. Other important metabolite classes, such as lipids, neurotransmitters, and proteins, which may also play key roles in AD pathology, were not assessed. Additionally, the experimental design included four discrete age time points (12, 25, 50, and 85 weeks), which allowed us to investigate broad age-related trends. However, because this study did not include intermediate time points (e.g., 18, 35, or 70 weeks), it may have overlooked transient evolving metabolic alterations that occur between the sampled stages. Such intermediate changes could be critical for identifying early biomarkers of metabolic disruptions during AD development. A more frequent sampling strategy might offer finer temporal resolution and additional insights into dynamic metabolic transitions during disease progression.

### Implications and Future Directions

Our findings have important implications for both mechanistic and translational AD research. The observed amine level shifts with age, sex and AD, along with their altered plasma-CSF correlation patterns, suggest that dysregulation in systemic amine metabolism, which may reflect changes in neurotransmission, inflammation, and oxidative stress, could contribute to the progression of AD pathology. Given the parallels between our findings in TgF344-AD rats and prior human studies, this AD rat model serves as a valuable platform for investigating AD-related processes. For example, Basun et al [16] reported significantly decreased plasma levels of glutamate and taurine, and increased CSF levels of glycine, leucine, and valine in AD patients compared to healthy controls. These human findings closely mirror our results in TgF344-AD rats, supporting the translational relevance of the model.

Additionally, several amines showed strong correlations between CSF and plasma, highlighting the potential for developing plasma-based biomarkers for AD. While we did not directly assess BBB transporter expression or activity in this study, the observed amine plasma-CSF associations for specific amines, such as 2-aminoadipic acid, kynurenine, and taurine, suggest that altered transporter function of relevant amino acid transport systems (e.g., system L, y+) may play a role in modulating amine flux across the BBB [58]. Future research should focus on validating these findings in human cohorts and systematically investigate the involvement of transporter proteins and metabolic enzymes regulating the homeostasis of amines, including but not limited to amino acids, in AD pathology.

Finally, the distinct sex differences observed in our study emphasize the need for sex-specific approaches in AD research and treatment. Understanding how AD related metabolic pathways differ between males and females could inform the development of personalized therapeutic strategies targeting disease progression at different stages [59].

## Conclusions

Overall, our findings provide novel insights into age-, sex-, and disease-dependent amine alterations in plasma and CSF in the TgF344-AD rat model. These results underscore amine dysregulation in AD and highlight the importance of considering both biological sex and disease stage in future mechanistic and biomarker studies. The TgF344-AD rat model proves to be a powerful tool for translational research, enabling the identification of metabolic disruptions and therapeutic targets relevant to AD progression.

## List of abbreviations

AD: Alzheimer’s disease
CSF: Cerebrospinal fluid
WT: Wildtype
GABA: Gamma-aminobutyric acid
BCAAs: Branched-chain amino acids
EDTA: Ethylenediaminetetraacetic acid
BMFL: Biomedical Metabolomics Facility Leiden
MRM: Multiple Reaction Monitoring
FDR: False discovery rate
GLM: Generalized logistic regression models
BBB: Blood-brain barrier
LAT: L-type amino acid transporter

## Acknowledgments

Chunyuan Yin acknowledges support from the China Scholarship Council (No. 202006550003). We thank the Predictive Pharmacology group for financing and executing the animal work and providing the samples. We thank the Biomedical Metabolomics Facility Leiden (BMFL) for their assistance with sample preparation and sample measurement.

## Ethics approval and consent to participate

Animal protocols were approved by the Leiden University Animal Welfare Body (AWB: AVD1060020171766). All animal experiments were conducted in accordance with the ARRIVE guidelines (ARRIVE 2.0).

## Funding

This publication is part of the “Building the infrastructure for Exposome research: Exposome-Scan” project (with project number 175.2019.032) of the program “Investment Grant NWO Large”, which is funded by the Dutch Research Council (NWO). This research was (partially) funded by X-Omics (NWO, project 184.034.019).

## Notes

### Competing Interest Statement

The authors have declared no competing interest.

## References

1. 2024 Alzheimer’s disease facts and figures. Alzheimers Dement, 2024. 20(5): p. 3708–3821.

2. Hardy, J. and D.J. Selkoe, The amyloid hypothesis of Alzheimer’s disease: progress and problems on the road to therapeutics. Science, 2002. 297(5580): p. 353–6.

3. Jack, C.R., Jr., et al., NIA-AA Research Framework: Toward a biological definition of Alzheimer’s disease. Alzheimers Dement, 2018. 14(4): p. 535–562.

4. Braak, H. and E. Braak, Neuropathological stageing of Alzheimer-related changes. Acta Neuropathol, 1991. 82(4): p. 239–59.

5. Release, B.P. FDA Approval of Aduhelm. 2021; Available from: https://investors.biogen.com/news-releases/news-release-details/fda-grants-accelerated-approval-aduhelmtm-first-and-only#:∼:text=(Tokyo%2C%20Japan)%20today%20announced,beta%20plaques%20in%20the%20brain.

6. Eisai/Biogen. FDA Approval of Leqembi. 2023; Available from: https://investors.biogen.com/news-releases/news-release-details/fda-advisory-committee-votes-unanimously-confirm-clinical#:∼:text=Lecanemab%20is%20a%20humanized%20immunoglobulin,)%20on%20January%206%2C%202023.

7. Meldrum, B., Amino acid neurotransmitters and new approaches to anticonvulsant drug action. Epilepsia, 1984. 25 Suppl 2: p. S140–9.

8. Smith, C.M., Marks’ basic medical biochemistry: A clinical approach. 2005: Lippincott Williams and Wilkins.

9. Purves, D., et al., Neurosciences. 2019: De Boeck Supérieur.

10. Socha, E., M. Koba, and P. Kośliński, Amino acid profiling as a method of discovering biomarkers for diagnosis of neurodegenerative diseases. Amino Acids, 2019. 51(3): p. 367–371.

11. Syeda, T. and J.R. Cannon, Potential Role of Heterocyclic Aromatic Amines in Neurodegeneration. Chem Res Toxicol, 2022. 35(1): p. 59–72.

12. Griffin, J.W. and P.C. Bradshaw, Amino acid catabolism in Alzheimer’s disease brain: friend or foe? Oxidative medicine and cellular longevity, 2017. 2017(1): p. 5472792.

13. Butterfield, D.A. and B. Halliwell, Oxidative stress, dysfunctional glucose metabolism and Alzheimer disease. Nature Reviews Neuroscience, 2019. 20(3): p. 148–160.

14. Ben-Ari, Y., et al., GABA: a pioneer transmitter that excites immature neurons and generates primitive oscillations. Physiol Rev, 2007. 87(4): p. 1215–84.

15. Yoo, H.S., U. Shanmugalingam, and P.D. Smith, Potential roles of branched-chain amino acids in neurodegeneration. Nutrition, 2022. 103-104: p. 111762.

16. Basun, H., et al., Amino acid concentrations in cerebrospinal fluid and plasma in Alzheimer’s disease and healthy control subjects. Journal of Neural Transmission-Parkinson’s Disease and Dementia Section, 1990. 2: p. 295–304.

17. Piubelli, L., et al., Serum D-serine levels are altered in early phases of Alzheimer’s disease: towards a precocious biomarker. Transl Psychiatry, 2021. 11(1): p. 77.

18. Kaiser, E., et al., Cerebrospinal fluid concentrations of functionally important amino acids and metabolic compounds in patients with mild cognitive impairment and Alzheimer’s disease. Neurodegener Dis, 2010. 7(4): p. 251–9.

19. Lim, N.K., et al., An Improved Method for Collection of Cerebrospinal Fluid from Anesthetized Mice. J Vis Exp, 2018(133).

20. Trushina, E., et al., Identification of altered metabolic pathways in plasma and CSF in mild cognitive impairment and Alzheimer’s disease using metabolomics. PLoS One, 2013. 8(5): p. e63644.

21. Molina, J.A., et al., Cerebrospinal fluid levels of non-neurotransmitter amino acids in patients with Alzheimer’s disease. Journal of Neural Transmission, 1998. 105(2): p. 279–286.

22. Yin, C., et al., Status of Metabolomic Measurement for Insights in Alzheimer’s Disease Progression-What Is Missing? Int J Mol Sci, 2023. 24(5).

23. Karu, N., et al., Severe COVID-19 Is Characterised by Perturbations in Plasma Amines Correlated with Immune Response Markers, and Linked to Inflammation and Oxidative Stress. Metabolites, 2022. 12(7).

24. van der Peet, M., et al., mzQuality: A tool for quality monitoring and reporting of targeted mass spectrometry measurements. bioRxiv, 2025: p. 2025.01. 22.633547.

25. Lyssikatos, C., et al., γ-Aminobutyric acids (GABA) and serum GABA/AABA (G/A) ratio as potential biomarkers of physical performance and aging. Scientific Reports, 2023. 13(1): p. 17083.

26. Zhong, L., et al., Key genes and pathways in asparagine metabolism in Alzheimer’s Disease: a bioinformatics approach. bioRxiv, 2025: p. 2025.04. 25.650586.

27. Brouquisse, R., et al., Asparagine metabolism and nitrogen distribution during protein degradation in sugar-starved maize root tips. Planta, 1992. 188(3): p. 384–395.

28. Penttinen, A., et al., Prolyl oligopeptidase: a rising star on the stage of neuroinflammation research. CNS & Neurological Disorders-Drug Targets (Formerly Current Drug Targets-CNS & Neurological Disorders), 2011. 10(3): p. 340–348.

29. Villa, A., et al., Estrogens, Neuroinflammation, and Neurodegeneration. Endocr Rev, 2016. 37(4): p. 372–402.

30. Chandran, S. and D. Binninger, Role of Oxidative Stress, Methionine Oxidation and Methionine Sulfoxide Reductases (MSR) in Alzheimer’s Disease. Antioxidants, 2023. 13(1): p. 21.

31. Moskovitz, J., et al., Induction of methionine-sulfoxide reductases protects neurons from amyloid β-protein insults in vitro and in vivo. Biochemistry, 2011. 50(49): p. 10687–10697.

32. Buccellato, F.R., et al., Role of Oxidative Damage in Alzheimer’s Disease and Neurodegeneration: From Pathogenic Mechanisms to Biomarker Discovery. Antioxidants (Basel), 2021. 10(9).

33. Sultana, R. and D.A. Butterfield, Role of oxidative stress in the progression of Alzheimer’s disease. J Alzheimers Dis, 2010. 19(1): p. 341–53.

34. Schöneich, C., Methionine oxidation by reactive oxygen species: reaction mechanisms and relevance to Alzheimer’s disease. Biochimica et Biophysica Acta (BBA)-Proteins and Proteomics, 2005. 1703(2): p. 111–119.

35. Vitvitsky, V., et al., A Functional Transsulfuration Pathway in the Brain Links to Glutathione Homeostasis *. Journal of Biological Chemistry, 2006. 281(47): p. 35785–35793.

36. Flori, L., et al., Transsulfuration Pathway Products and H2S-Donors in Hyperhomocysteinemia: Potential Strategies Beyond Folic Acid. International Journal of Molecular Sciences, 2025. 26(13): p. 6430.

37. Dumont, M. and M.F. Beal, Neuroprotective strategies involving ROS in Alzheimer disease. Free Radic Biol Med, 2011. 51(5): p. 1014–26.

38. Granot-Hershkovitz, E., et al., Genetic loci of beta-aminoisobutyric acid are associated with aging-related mild cognitive impairment. Translational Psychiatry, 2023. 13(1): p. 140.

39. Mahajan, U.V., et al., Dysregulation of multiple metabolic networks related to brain transmethylation and polyamine pathways in Alzheimer disease: A targeted metabolomic and transcriptomic study. PLoS medicine, 2020. 17(1): p. e1003012.

40. Ozaki, T., et al., Metabolomic alterations in the blood plasma of older adults with mild cognitive impairment and Alzheimer’s disease (from the Nakayama Study). Scientific Reports, 2022. 12(1): p. 15205.

41. Polis, B., D. Karasik, and A.O. Samson, Alzheimer’s disease as a chronic maladaptive polyamine stress response. Aging (Albany NY), 2021. 13(7): p. 10770.

42. El-Halfawy, O.M. and M.A. Valvano, Putrescine reduces antibiotic-induced oxidative stress as a mechanism of modulation of antibiotic resistance in Burkholderia cenocepacia. Antimicrobial agents and chemotherapy, 2014. 58(7): p. 4162–4171.

43. Arthur, R., S. Jamwal, and P. Kumar, A review on polyamines as promising nextgeneration neuroprotective and anti-aging therapy. European Journal of Pharmacology, 2024: p. 176804.

44. Makletsova, M., et al., The role of polyamines in the mechanisms of cognitive impairment. Neurochemical Journal, 2022. 16(3): p. 283–294.

45. Dayon, L., et al., Proteomes of paired human cerebrospinal fluid and plasma: relation to blood–brain barrier permeability in older adults. Journal of proteome research, 2019. 18(3): p. 1162–1174.

46. Bruno, M., et al., Blood–brain barrier permeability is associated with different neuroinflammatory profiles in Alzheimer’s disease. European journal of neurology, 2024. 31(1): p. e16095.

47. Chalbot, S., et al., Blood-cerebrospinal fluid barrier permeability in Alzheimer’s disease. Journal of Alzheimer’s Disease, 2011. 25(3): p. 505–515.

48. Ott, B.R., et al., Blood-cerebrospinal fluid barrier gradients in mild cognitive impairment and Alzheimer’s disease: relationship to inflammatory cytokines and chemokines. Frontiers in aging neuroscience, 2018. 10: p. 245.

49. Harder, A.V., et al., Correlation of Amine Concentrations in Blood and Cerebrospinal Fluid in Healthy Volunteers and Migraineurs. International Journal of Molecular Sciences, 2025. 26(20): p. 9899.

50. Smith, Q.R., Transport of glutamate and other amino acids at the blood-brain barrier. The Journal of nutrition, 2000. 130(4): p. 1016S–1022S.

51. Liang, Y., et al., Kynurenine pathway metabolites as biomarkers in Alzheimer’s disease. Disease Markers, 2022. 2022(1): p. 9484217.

52. Sharma, V.K., et al., Kynurenine metabolism and Alzheimer’s disease: the potential targets and approaches. Neurochemical Research, 2022. 47(6): p. 1459–1476.

53. Kearns, R., Gut–brain axis and neuroinflammation: The role of gut permeability and the kynurenine pathway in neurological disorders. Cellular and Molecular Neurobiology, 2024. 44(1): p. 64.

54. Fukui, S., et al., Blood–brain barrier transport of kynurenines: implications for brain synthesis and metabolism. Journal of neurochemistry, 1991. 56(6): p. 2007–2017.

55. Huf, F., et al., Neuroprotection elicited by taurine in sporadic Alzheimer-like disease: benefits on memory and control of neuroinflammation in the hippocampus of rats. Molecular and Cellular Biochemistry, 2024. 479(10): p. 2663–2678.

56. Rafiee, Z., A.M. García-Serrano, and J.M. Duarte, Taurine supplementation as a neuroprotective strategy upon brain dysfunction in metabolic syndrome and diabetes. Nutrients, 2022. 14(6): p. 1292.

57. Kang, Y.S., et al., Regulation of taurine transport at the blood–brain barrier by tumor necrosis factor-α, taurine and hypertonicity. Journal of neurochemistry, 2002. 83(5): p. 1188–1195.

58. Baloni, P., et al., Metabolic Network Analysis Reveals Altered Bile Acid Synthesis and Metabolism in Alzheimer’s Disease. Cell Rep Med, 2020. 1(8): p. 100138.

59. Robison, L.S., et al., Sex differences in metabolic phenotype and hypothalamic inflammation in the 3xTg-AD mouse model of Alzheimer’s disease. Biol Sex Differ, 2023. 14(1): p. 51.

