## Supplementary material for "Longitudinal Metabolomic Profiling of Biogenic Amines in Plasma and CSF, and Their Correlation, Reveals Sex-Specific and Age Changes in TgF344 Alzheimer’s Disease Transgenic and Wildtype Rats": Supplementary Tables 3-9.docx

To provide a more detailed overview of the metabolite-specific plasma-CSF correlation patterns, supplementary Tables 3-9 summarize the pearson correlation coefficients for each metabolite across TgF344-AD, WT, and combined (ADWT) groups, grouped by sex (Supplementary Tables 4-5) and age (Supplementary Tables 6-9). For each group, the Fisher’s z-transformed correlation coefficients, corresponding p-values, and FDR-adjusted q-values are provided. Only metabolites that showed a statistically supported correlation (|R| > 0.35, *p* < 0.05) in at least one group (e.g., TgF344-AD, WT, or ADWT, stratified by age or sex) were included in the supplementary table. Results are rounded to two decimal places, and non-significant comparisons (*p* > 0.05) are labeled as “n.s.”

Supplementary Table-3: The correlation coefficient between CSF and plasma of the metabolites within AD, WT or a combination of AD and WT (ADWT) and the fisher's transformed z-score of the AD correlation coefficient (n = 136) and WT correlation coefficient (n = 128) for all rats. Only the metabolites that have at least one significant (^*^*p*<0.05) correlation coefficient higher than |0.35| in AD, WT and/or ADWT are displayed in the table. It is rounded in two digits. The *p*-value and FDR adjusted *p*-value (*q*-value) are calculated with a t-test. (^ns^*p*≥0.05)

|  | Correlation coefficient | | | Fisher's Z-value AD/WT | | |
| --- | --- | --- | --- | --- | --- | --- |
| Amine | R_AD_ | R_WT_ | R_(ADWT)_ | z-value | *p*-value | *q*-value |
| 1-methylhistidine | 0.52 | 0.49 | 0.51 | 0.27 | 7.86e-01 | 0.82 |
| 2-aminoadipic acid | 0.38 | 0.09^ns^ | 0.24 | 2.48 | 1.32e-02 | 0.06 |
| 3-methoxytyrosine | 0.35 | 0.21^ns^ | 0.28 | 1.20 | 2.32e-01 | 0.39 |
| Alpha aminobutyric acid | 0.80 | 0.46 | 0.63 | 4.91 | 0 | 0 |
| Citrulline | -0.68 | -0.52 | -0.60 | -2.04 | 4.16e-02 | 0.14 |
| DL-3-aminoisobutyric acid | 0.42 | -0.19^ns^ | 0.17^ns^ | 5.12 | 0 | 0 |
| Kynurenine | 0.59 | 0.20^ns^ | 0.45 | 3.76 | 2.0e-4 | 0.003 |
| Methionine sulfoxide | -0.37 | -0.27 | -0.32 | -0.91 | 3.64e-01 | 0.49 |
| N6,N6,N6-trimethyl lysine | 0.60 | 0.40 | 0.52 | 2.10 | 3.61e-02 | 0.13 |
| Phenylalanine | 0.41 | 0.07^ns^ | 0.26 | 2.91 | 3.6e-03 | 0.03 |
| Putrescine | 0.60 | 0.20^ns^ | 0.42 | 3.91 | 1.0e-04 | 0.002 |
| Taurine | 0.47 | 0.28 | 0.39 | 1.72 | 8.46e-02 | 0.22 |

Supplementary Table-4: The correlation coefficient between CSF and plasma of the metabolites within AD, WT or a combination of AD and WT (ADWT) and the fisher's transformed z-score of the AD correlation coefficient (n = 62) and WT correlation coefficient (n = 72) for the male rats. Only the metabolites that have at least one significant (^*^*p*<0.05) correlation coefficient higher than |0.35| in AD, WT and/or ADWT are displayed in the table. It is rounded in two digits. The p-value and FDR adjusted p-value (*q-*value) are calculated with a t-test. (^ns^*p*≥0.05)

|  | Correlation coefficient | | | Fisher's Z-value AD/WT | | |
| --- | --- | --- | --- | --- | --- | --- |
| Amine | R_AD_ | R_WT_ | R_(ADWT)_ | z-value | *p*-value | *q*-value |
| 1-methylhistidine | 0.44 | 0.37 | 0.42 | 0.53 | 6.0e-01 | 0.79 |
| 2-aminoadipic acid | 0.42 | -0.03^ns^ | 0.22^ns^ | 2.68 | 7.3e-03 | 0.06 |
| Alpha-aminobutyric acid | 0.79 | 0.54 | 0.67 | 2.72 | 6.6e-03 | 0.06 |
| Citrulline | -0.69 | -0.63 | -0.65 | -0.57 | 5.70e-01 | 0.76 |
| DL-3-aminoisobutyric acid | 0.49 | -0.32^ns^ | 0.14^ns^ | 4.84 | 0 | 0 |
| Gamma-aminobutyric acid | 0.06^ns^ | -0.39 | -0.18^ns^ | 2.653 | 0.008 | 0.06 |
| Gamma glutamyl alanine | 0.03^ns^ | 0.51 | 0.26 | -3.03 | 2.4e-03 | 0.04 |
| Kynurenine | 0.64 | 0.31^ns^ | 0.53 | 2.46 | 1.39e-02 | 0.06 |
| Leucine | 0.04^ns^ | -0.4 | -0.13^ns^ | 2.62 | 8.9e-03 | 0.06 |
| Methionine sulfone | -0.02^ns^ | 0.4 | 0.16^ns^ | -2.52 | 1.19e-02 | 0.06 |
| Methionine.sulfoxide | -0.39 | -0.34 | -0.37 | -0.35 | 7.28e-01 | 0.83 |
| N6,N6,N6-trimethyl lysine | 0.53 | 0.54 | 0.54 | -0.11 | 9.13e-01 | 0.93 |
| Putrescine | 0.49 | 0.22^ns^ | 0.35 | 1.74 | 8.13e-02 | 0.22 |
| Saccharopine | -0.18^ns^ | 0.42 | 0.08^ns^ | -3.57 | 4.0e-04 | 0.01 |
| Sarcosine | -0.06^ns^ | -0.48 | -0.24 | 2.57 | 1.01e-02 | 0.06 |
| Taurine | 0.54 | 0.17^ns^ | 0.36 | 2.46 | 1.4e-02 | 0.06 |

Supplementary Table-5: The correlation coefficient between CSF and plasma of the metabolites within AD, WT or a combination of AD and WT (ADWT) and the fisher's transformed z-score of the AD correlation coefficient (n = 74) and WT correlation coefficient (n = 56) for the female rats. Only the metabolites that have at least one significant (^*^*p*<0.05) correlation coefficient higher than |0.35| in AD, WT and/or ADWT are displayed in the table. It is rounded in two digits. The *p*-value and FDR adjusted p-value (*q*-value) are calculated with a t-test. (^ns^*p*≥0.05)

|  | Correlation coefficient | | | Fisher's Z-value AD/WT | | |
| --- | --- | --- | --- | --- | --- | --- |
| Amine | R_AD_ | R_WT_ | R_(ADWT)_ | z-value | *p*-value | *q*-value |
| 1-methylhistidine | 0.68 | 0.51 | 0.55 | 1.45 | 1.47e-01 | 0.30 |
| 2-aminoadipic acid | 0.36 | 0.31^ns^ | 0.29 | 0.31 | 7.57e-01 | 0.84 |
| Alpha-aminobutyric acid | 0.70 | 0.09^ns^ | 0.37 | 4.31 | 0 | 0 |
| Citrulline | -0.68 | -0.38 | -0.53 | -2.34 | 1.91e-02 | 0.08 |
| Cysteine | 0.04^ns^ | -0.38 | -0.21^ns^ | 2.45 | 1.43e-02 | 0.08 |
| DL-3-aminoisobutyric acid | 0.35 | 0.05^ns^ | 0.23^ns^ | 1.73 | 8.46e-02 | 0.22 |
| Glycylglycine | 0.39 | -0.02^ns^ | 0.23^ns^ | 2.39 | 1.71e-02 | 0.08 |
| Kynurenine | 0.53 | 0.12^ns^ | 0.40 | 2.56 | 1.06e-02 | 0.08 |
| Methionine sulfoxide | -0.36 | -0.18^ns^ | -0.28 | -1.10 | 2.70e-01 | 0.49 |
| N6,N6,N6-trimethyl lysine | 0.64 | 0.25^ns^ | 0.50 | 2.77 | 5.6e-03 | 0.05 |
| O-acetyl-serine | 0.08^ns^ | -0.42 | -0.17^ns^ | 2.94 | 3.3e-03 | 0.03 |
| Phenylalanine | 0.48 | 0.09^ns^ | 0.31 | 2.40 | 1.66e-02 | 0.08 |
| Putrescine | 0.67 | 0.18^ns^ | 0.46 | 3.49 | 5.0e-04 | 0.009 |
| Serotonine | 0.09^ns^ | 0.38 | 0.22^ns^ | -1.70 | 8.87e-02 | 0.22 |
| S-methylcysteine | 0.49 | -0.35^ns^ | 0.06^ns^ | 5.02 | 0 | 0 |
| Taurine | 0.42 | 0.56 | 0.47 | -1.06 | 2.88e-01 | 0.51 |

Supplementary Table-6 The correlation coefficient between CSF and plasma of the metabolites within AD, WT or a combination of AD and WT (ADWT) and the fisher's transformed z-score of the AD correlation coefficient (n= 26) and WT correlation coefficient (n = 24) for the 12 weeks old rats. Only the metabolites that have at least one significant (^*^*p*<0.05) correlation coefficient higher than |0.35| in AD, WT and/or ADWT are displayed in the table. It is rounded in two digits. The p-value and FDR adjusted p-value (*q*-value) are calculated with a t-test. (^ns^*p*≥0.05)

|  | Correlation coefficient | | | Fisher's Z-value AD/WT | | |
| --- | --- | --- | --- | --- | --- | --- |
| Amine | R_AD_ | R_WT_ | R_(ADWT)_ | z-value | *p*-value | *q*-value |
| 1-methylhistidine | 0.70 | 0.67 | 0.67 | 0.14 | 8.913e-01 | 0.92 |
| Alpha-aminobutyric acid | 0.84 | 0.42^ns^ | 0.64 | 2.51 | 1.21e-02 | 0.04 |
| Asparagine | 0.38^ns^ | -0.66 | -0.08^ns^ | 3.96 | 1.0e-04 | 0.001 |
| Citrulline | -0.81 | -0.80 | -0.76 | -0.06 | 9.51e-01 | 0.951 |
| Cysteine | 0.32^ns^ | -0.60 | -0.22^ns^ | 3.39 | 7.0e-04 | 0.004 |
| DL-3-aminobutyric acid | 0.74 | -0.21^ns^ | 0.21^ns^ | 3.91 | 1.0e-04 | 0.001 |
| Glutamine | -0.15^ns^ | -0.80 | -0.34^ns^ | 3.08 | 2.1e-03 | 0.01 |
| Glycylglycine | 0.76 | -0.34^ns^ | 0.27^ns^ | 4.46 | 0 | 0 |
| Hydroxylysine | -0.54^ns^ | -0.63 | -0.58 | 0.44 | 6.613e-01 | 0.76 |
| Kynurenine | 0.61 | 0.3^ns^ | 0.51 | 1.32 | 1.867e-01 | 0.32 |
| Methionine sulfoxide | -0.43^ns^ | -0.66 | -0.52 | 1.11 | 2.654e-01 | 0.36 |
| N6,N6,N6-trimethyl lysine | 0.55^ns^ | 0.57^ns^ | 0.56 | -0.12 | 9.07e-01 | 0.92 |
| Norepinephrine | -0.32^ns^ | -0.68 | -0.36^ns^ | 1.63 | 1.04e-01 | 0.20 |
| Phenylalanine | 0.69 | -0.31^ns^ | 0.29^ns^ | 3.88 | 1.0e-04 | 0.001 |
| Proline | 0.15^ns^ | -0.84 | -0.42 | 4.60 | 0 | 0 |
| Putrescine | 0.66 | 0.39^ns^ | 0.52 | 1.26 | 2.061e-01 | 0.33 |
| Sarcosine | -0.13^ns^ | -0.66 | -0.44 | 2.20 | 2.75e-02 | 0.08 |
| S-methylcysteine | 0.11^ns^ | -0.58 | -0.15^ns^ | 2.57 | 1.01e-02 | 0.04 |
| Taurine | 0.78 | 0.26^ns^ | 0.57 | 2.59 | 9.6e-03 | 0.04 |

Supplementary Table-7: The correlation coefficient between CSF and plasma of the metabolites within AD, WT or a combination of AD and WT (ADWT) and the fisher's transformed z-score of the correlation coefficient of AD (n = 38) and the correlation coefficient of WT (n = 36) for the 25 weeks old rats. Only the metabolites that have at least one significant (^*^*p*<0.05) correlation coefficient higher than |0.35| in AD, WT and/or ADWT are displayed in the table. It is rounded in two digits. The p-value and FDR adjusted p-value (*q*-value) are calculated with a t-test. (^ns^*p*≥0.05)

|  | Correlation coefficient | | | Fisher's Z-value AD/WT | | |
| --- | --- | --- | --- | --- | --- | --- |
| Amine | R_AD_ | R_WT_ | R_(ADWT)_ | z-value | *p*-value | *q*-value |
| 1-methylhistidine | 0.74 | 0.11^ns^ | 0.45 | 3.41 | 7.0e-04 | 0.02 |
| Alpha-aminobutyric acid | 0.69 | 0.88 | 0.82 | -2.17 | 3.03e-02 | 0.15 |
| Citrulline | -0.38^ns^ | -0.84 | -0.64 | 3.33 | 9.0e-04 | 0.02 |
| Glycylglycine | 0.14^ns^ | -0.62 | -0.10^ns^ | 3.56 | 4.0e-04 | 0.02 |
| Hydroxylysine | -0.18^ns^ | -0.47 | -0.25^ns^ | 1.35 | 1.76e-01 | 0.41 |
| Leucine | 0.03^ns^ | -0.50 | -0.25^ns^ | 2.39 | 1.69e-02 | 0.13 |
| Methionine | 0.46 | 0.23^ns^ | 0.18^ns^ | 1.07 | 2.83e-01 | 0.49 |
| N6,N6,N6-trimethyl lysine | 0.59 | 0.42^ns^ | 0.49 | 0.92 | 3.57e-01 | 0.56 |
| Proline | -0.06^ns^ | -0.53 | -0.39 | 2.18 | 2.91e-02 | 0.15 |
| Sarcosine | -0.10^ns^ | -0.56 | -0.32^ns^ | 2.23 | 2.58e-02 | 0.15 |

Supplementary Table-8: The correlation coefficient between CSF and plasma of the metabolites within AD, WT or a combination of AD and WT (ADWT) and the fisher's transformed z-score of the correlation coefficient in AD (n = 36) and the correlation coefficient in WT (n = 32) for the 50 weeks old rats. Only the metabolites that have at least one significant (^*^*p*<0.05) correlation coefficient higher than |0.35| in AD, WT and/or ADWT are displayed in the table. It is rounded in two digits. The p-value and FDR adjusted p-value (*q*-value) are calculated with a t-test. (^ns^*p*≥0.05)

|  | Correlation coefficient | | | Fisher's Z-value AD/WT | | |
| --- | --- | --- | --- | --- | --- | --- |
| Amine | R_AD_ | R_WT_ | R_(ADWT)_ | z-value | *p*-value | *q*-value |
| 1-methylhistidine | 0.17^ns^ | 0.58 | 0.31^ns^ | -1.92 | 5.51e-02 | 0.32 |
| 3-methoxytyrosine | 0.39^ns^ | 0.36^ns^ | 0.41 | 0.11 | 9.12e-01 | 0.97 |
| Alpha-aminobutyric acid | 0.73 | 0.70 | 0.71 | 0.29 | 7.74e-01 | 0.97 |
| Citrulline | -0.82 | -0.47^ns^ | -0.66 | -2.54 | 1.1e-02 | 0.12 |
| DL-3-aminoisobutyric acid | 0.58 | -0.06^ns^ | 0.32^ns^ | 2.86 | 4.3e-03 | 0.08 |
| Kynurenine | 0.88 | 0.18^ns^ | 0.61 | 4.70 | 0 | 0 |
| Methionine sulfone | 0.24^ns^ | 0.32^ns^ | 0.35 | -0.38 | 7.02e-01 | 0.97 |
| Methionine sulfoxide | -0.44^ns^ | -0.27^ns^ | -0.37 | -0.73 | 4.68e-01 | 0.80 |
| N6,N6,N6-trimethyl lysine | 0.69 | 0.34^ns^ | 0.53 | 1.93 | 5.39e-02 | 0.32 |
| Phenylalanine | 0.52 | -0.13^ns^ | 0.25^ns^ | 2.76 | 5.7e-03 | 0.08 |
| Putrescine | 0.77 | 0.02^ns^ | 0.44 | 3.91 | 1.0e-04 | 0.003 |

Supplementary Table-9: The correlation coefficient between CSF and plasma of the metabolites within AD, WT or a combination of AD and WT (ADWT) and the fisher's transformed z-score of the correlation coefficient in AD (n = 36) and the correlation coefficient in WT (n = 36) for the 85 weeks old rats. Only the metabolites that have at least one significant (^*^*p*<0.05) correlation coefficient higher than |0.35| in AD, WT and/or ADWT are displayed in the table. It is rounded in two digits. The p-value and FDR adjusted p-value (*q*-value) are calculated with a t-test. (^ns^*p*≥0.05)

|  | Correlation coefficient | | | Fisher's Z-value AD/WT | | |
| --- | --- | --- | --- | --- | --- | --- |
| Amine | R_AD_ | R_WT_ | R_(ADWT)_ | z-value | *p*-value | *q*-value |
| 1-methylhistidine | 0.56 | 0.63 | 0.60 | -0.49 | 6.27e-01 | 0.77 |
| Alpha-aminobutyric acid | 0.89 | 0.13^ns^ | 0.43 | 5.30 | 0 | 0 |
| Anserine | 0.40^ns^ | 0.53 | 0.45 | -0.70 | 4.86e-01 | 0.77 |
| Citrulline | -0.63 | 0.08^ns^ | -0.27^ns^ | -3.31 | 9.0e-04 | 0.01 |
| Gamma glutamyl alanine | -0.40^ns^ | 0.53 | 0.16^ns^ | -4.12 | 0 | 0 |
| Glutamine | 0.48 | -0.19^ns^ | 0.23^ns^ | 2.94 | 3.3e-03 | 0.02 |
| Histidine | 0.20^ns^ | 0.54 | 0.40 | -1.63 | 1.02e-01 | 0.25 |
| Kynurenine | 0.54 | 0.1^ns^ | 0.33 | 2.06 | 3.95e-02 | 0.13 |
| Lysine | 0.29^ns^ | 0.42^ns^ | 0.37 | -0.59 | 5.55e-01 | 0.77 |
| N6,N6,N6-trimethyl lysine | 0.67 | 0.35 ^ns^ | 0.53 | 1.77 | 7.76e-02 | 0.20 |
| O-acetyl serine | 0.57 | -0.41^ns^ | 0.04^ns^ | 4.37 | 0 | 0 |
| Ornithine | 0.41^ns^ | 0.51 | 0.48 | -0.48 | 6.34e-01 | 0.77 |
| Phenylalanine | 0.31^ns^ | 0.57 | 0.43 | -1.32 | 1.88e-01 | 0.41 |
| Putrescine | 0.71 | 0.13 ^ns^ | 0.41 | 3.12 | 1.8e-03 | 0.02 |
| Saccharopine | -0.58 | 0.31^ns^ | -0.24^ns^ | -4.02 | 1.0e-04 | 0.001 |
| S-methylcysteine | 0.68 | 0.13^ns^ | 0.42 | 2.86 | 4.2e-03 | 0.03 |
| Taurine | 0.62 | 0.22^ns^ | 0.43 | 2.03 | 4.28e-02 | 0.13 |
| Tryptophan | 0.31^ns^ | 0.48 | 0.40 | -0.82 | 4.12e-01 | 0.70 |
| Tyrosine | 0.11^ns^ | 0.52 | 0.27^ns^ | -1.85 | 6.45e-02 | 0.17 |
| Valine | 0.22^ns^ | 0.45^ns^ | 0.36 | 1.03 | 3.04e-01 | 0.56 |
