## Supplementary figures and images for "Longitudinal Metabolomic Profiling of Biogenic Amines in Plasma and CSF, and Their Correlation, Reveals Sex-Specific and Age Changes in TgF344 Alzheimer’s Disease Transgenic and Wildtype Rats"

### Supplementary Figure 1.pdf

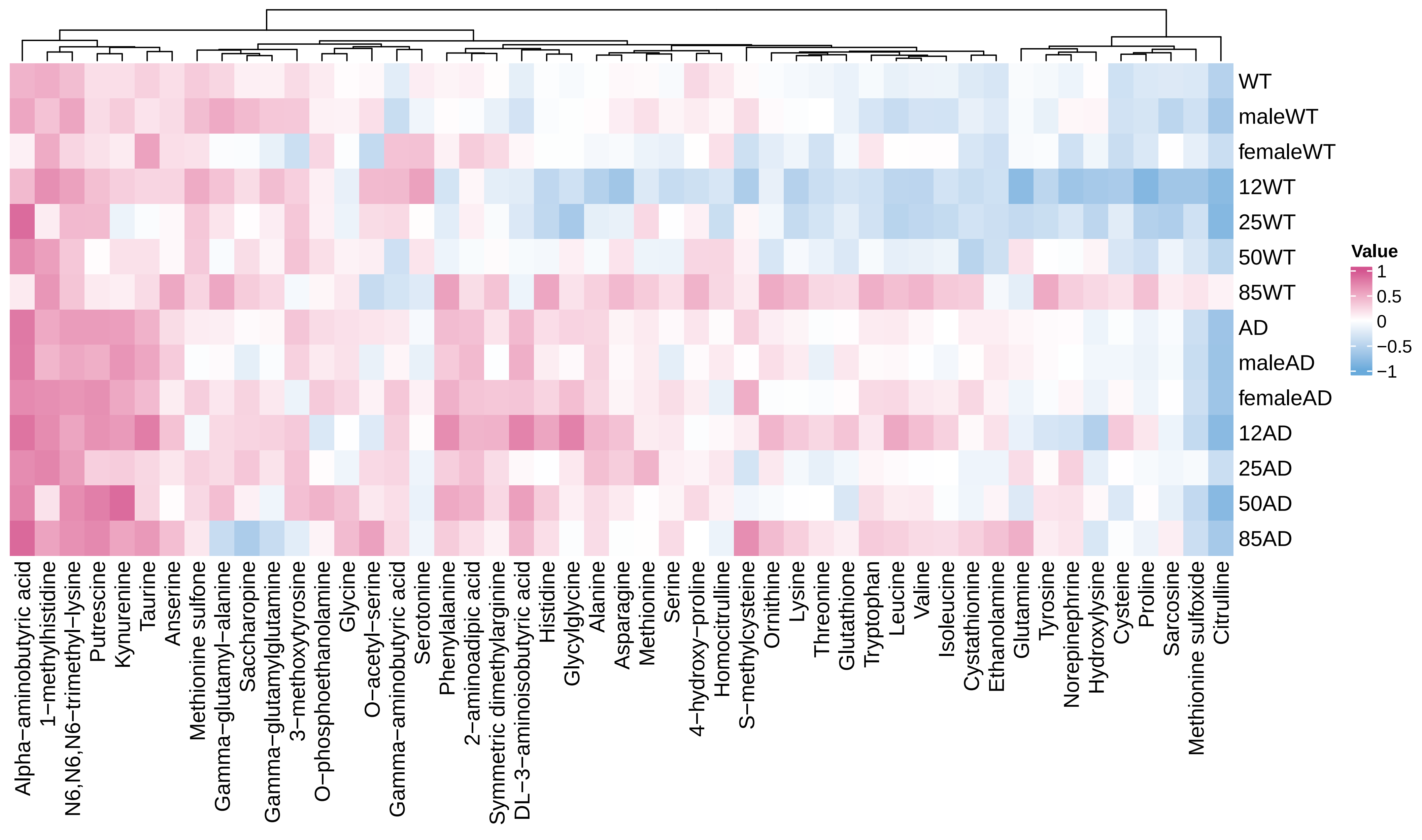

### Supplementary Figure 2.pdf

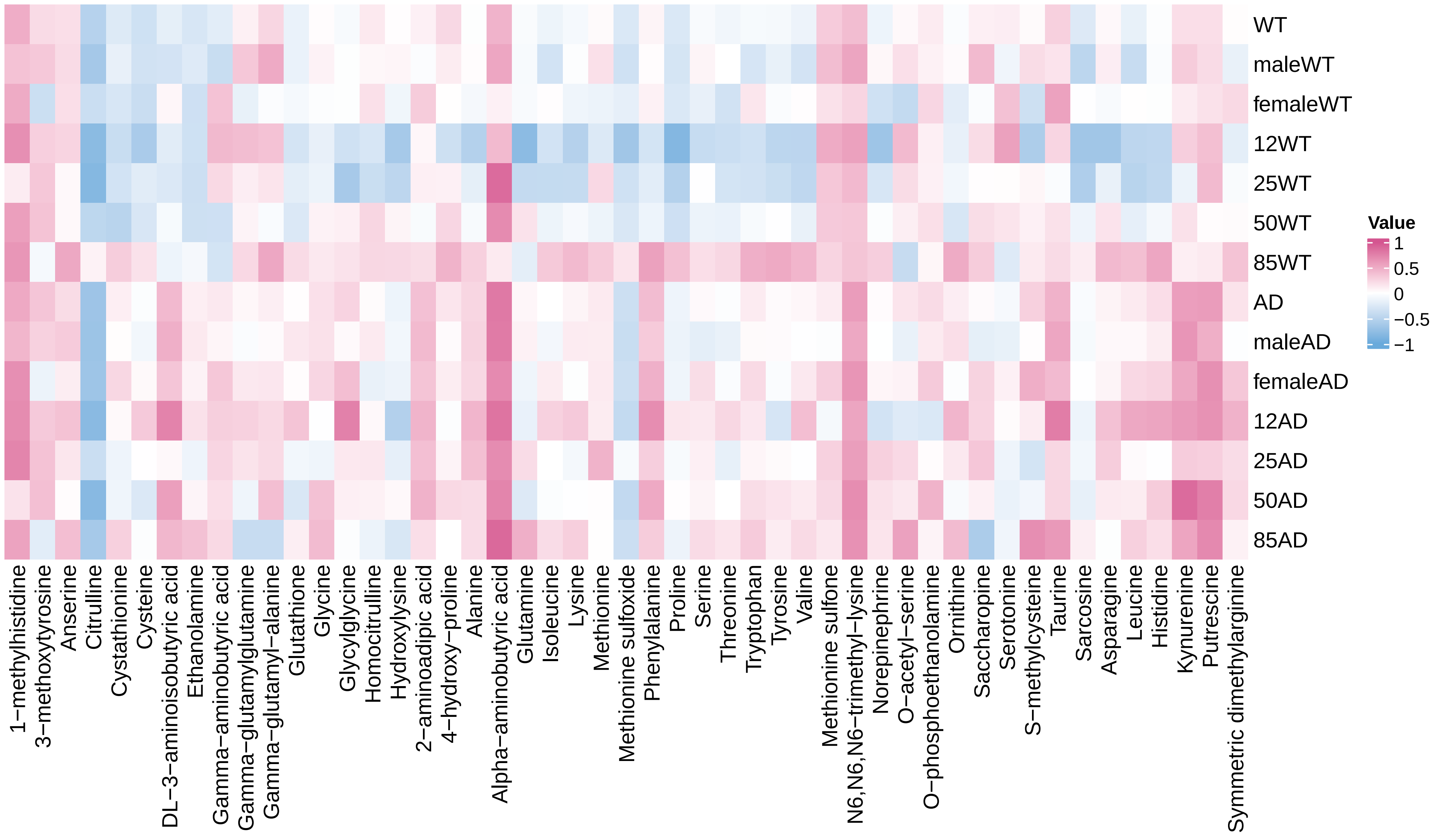
